# Dynamic Consistency Reveals Predictable Genes in Cross-Cell Type Temporal scRNA-Seq Data

**DOI:** 10.64898/2026.02.11.705300

**Authors:** Jiajie Shi, Rui Wu, Yushi Liu, Rui Li, Alexandre Duprey

## Abstract

Understanding how gene expression evolves over time after trauma is central to modeling immune responses, yet single-cell temporal data remain sparse and heterogeneous across cell types. Using a temporal trauma scRNA-seq dataset, we formulate the task of predicting next-time gene expression from earlier observations under a cross-cell-type generalization setting. We introduce the *Dynamic Consistency Index* (DCI), which quantifies how consistently a gene’s temporal trajectory aligns across cell types, serving as a measure of biological regularity and predictability. High-DCI genes exhibit reproducible temporal dynamics and are markedly easier to model. By integrating DCI-based gene selection with a recurrent neural architecture trained under a Gaussian negative log-likelihood objective, we achieve superior accuracy and well-calibrated uncertainty compared to deterministic baselines. Overall, DCI reliably identifies dynamically consistent genes, and uncertainty-aware recurrent modeling provides a robust framework for capturing cross-cell-type gene-expression evolution.

## Introduction

Modeling how gene expression evolves over time is central to understanding the biological response following perturbations such as trauma or infection. Advances in single-cell RNA sequencing (scRNA-seq) now allow measurements across multiple cell types and timepoints, providing an opportunity to study temporal regulation at cellular resolution. However, quantitative modeling of such dynamics remains challenging. Expression trajectories are often short, noisy, and heterogeneous across cell types, making it difficult to identify reproducible temporal patterns or to extrapolate behavior in unseen contexts.

Most existing approaches focus on reconstructing latent trajectories or transition maps from static snapshots, as in pseudotime inference or RNA velocity analysis (Bergen et al. 2020; Qiu et al. 2022). These techniques are effective for visualizing developmental flows but are not designed to predict future expression levels or quantify uncertainty. Furthermore, models trained within one cell type or lineage seldom generalize to others, limiting their ability to capture global regularities across heterogeneous populations.

Temporal prediction across cell types provides a rigorous test of whether a model captures generalizable biological programs rather than cell-type–specific fluctuations. Biologically, different immune cell populations sometimes share coordinated transcriptional responses to trauma—such as inflammation, stress signaling, and metabolic reprogramming—yet manifest them with distinct magnitudes and timings. Accurately predicting temporal gene-expression changes in one cell type based on patterns learned from others therefore reflects an understanding of these shared regulatory mechanisms.

In practice, however, many time-series models easily fit within a single cell type but fail to generalize to others. This is because that not all genes follow shared temporal programs—some exhibit consistent trajectories across cell types, while others behave idiosyncratically or even oppositely. This variability makes global prediction across cell types inherently difficult and highlights the need to distinguish predictable from unpredictable genes.

From a biological perspective, genes with coherent dynamics often participate in coordinated immune responses, reflecting conserved regulatory mechanisms that unfold similarly across lineages. In contrast, context-dependent or stochastic genes reflect specialized cellular roles or noise, offering limited generalization. Therefore, rather than attempting to model all genes uniformly, our goal is to identify those with consistent dynamics across cell types and develop predictive models for this subset.

Using data from (Chen et al. 2021), we study the problem of *cross-cell-type temporal prediction* of genes in a human trauma scRNA-seq dataset, where each gene’s mean expression is aggregated at the cell-type level over four timepoints.

### Cross-cell-type temporal prediction problem

In single-cell RNA-seq time-series, many cell types are unevenly sampled across conditions or timepoints, and rare populations (e.g., T-cell subtypes) are often missing at later stages. We therefore consider the problem:

> *Given the temporal dynamics of a gene observed in a subset of cell types, can we infer its dynamics in unseen cell types?*

This is especially valuable for human trauma or disease datasets, where collecting balanced longitudinal samples for every cell population is infeasible.

The task is inherently challenging because different genes exhibit distinct temporal behaviors across cell types. These heterogeneous dynamics—where some genes rise sharply while others remain stable or even invert direction—make it difficult to construct a single model that generalizes across the entire transcriptome.

To address this, we define the Dynamic Consistency Index (DCI), which serves as a diagnostic measure—revealing which genes exhibit reproducible temporal patterns and are thus amenable to modeling. This framework reframes cross-cell-type prediction not as a solved problem, but as a structured challenge of separating inherently predictable biological signals from context-specific variability. DCI is computed by measuring pairwise alignment of temporal difference vectors in log-expression space. Genes with high DCI display coherent temporal trends—rising or falling synchronously across cell types—while those with low DCI behave irregularly or remain static. This index effectively distinguishes dynamically consistent genes that are predictable from those dominated by noise. By filtering on DCI, we isolate genes whose time evolution reflects reproducible biological processes.

For modeling, we employ a recurrent neural network trained with a Gaussian negative log-likelihood (NLL) objective. The model takes as input the time-series summary statistics for each gene and outputs both the predicted mean and variance at the next timepoint. This heteroscedastic formulation allows the model to express uncertainty proportional to biological variability, stabilizing training and improving generalization. Compared to feed-forward and transformer baselines, the recurrent Gaussian model achieves lower mean absolute error (MAE) and produces well-calibrated uncertainty estimates, particularly for high-DCI genes.

We evaluate the method under a cross-cell-type setting: models are trained on one subset of cell types and tested on disjoint ones. The results show that temporal consistency, as captured by DCI, is a strong indicator of predictability. Genes with high DCI yield accurate and stable predictions, while low-DCI genes are intrinsically unpredictable regardless of model complexity. The combination of DCI-based selection and uncertainty-aware recurrent modeling provides a practical framework for studying gene-expression dynamics in heterogeneous single-cell systems.

Our contributions are threefold:

- We introduce the **Dynamic Consistency Index (DCI)**, a simple and interpretable measure that quantifies temporal regularity of gene-expression trajectories across cell types.
- We develop an **uncertainty-aware recurrent model** trained with a Gaussian NLL loss to jointly predict expression means and variances.
- We demonstrate that high-DCI genes exhibit predictable temporal behavior, and that the proposed recurrent model achieves superior accuracy and uncertainty calibration in cross-cell-type prediction.

Together, these components reveal that temporal consistency is a key property distinguishing predictable genes from those dominated by stochastic fluctuations, offering a new perspective for modeling and interpreting dynamic scRNA-seq data.

## Related Work

### Temporal modeling in single-cell transcriptomics

Several methods have been developed to infer dynamic trajectories from static scRNA-seq data. RNA velocity (La Manno et al. 2018) introduced a first-order model of transcriptional change by comparing unspliced and spliced mRNA counts, while scVelo (Bergen et al. 2020) extended this idea to transient cell states through a dynamical system formulation. Dynamo (Qiu et al. 2022) generalized velocity into continuous vector fields, enabling reconstruction of global transcriptomic flows. Other approaches such as Monocle 3 (Cao et al. 2019) and Palantir (Setty et al. 2019) recover pseudotemporal orderings of cells along differentiation paths. These frameworks focus on reconstructing latent developmental landscapes rather than quantitatively predicting future expression levels. In contrast, our work treats temporal progression as a supervised prediction problem and evaluates generalization across cell types.

### Uncertainty-aware regression and heteroscedastic modeling

Neural models that estimate predictive variance have been studied extensively in machine learning. Kendall and Gal (Kendall and Gal 2017) formulated heteroscedastic regression via a Gaussian negative log-likelihood loss, enabling joint estimation of mean and variance. Extensions have appeared in molecular property prediction (Scalia et al. 2020) and gene-expression modeling (Zhou et al. 2024). In biological contexts, uncertainty estimates are valuable for distinguishing systematic temporal trends from stochastic noise. Our recurrent Gaussian model follows this line by explicitly modeling cell-type-specific uncertainty in temporal gene-expression prediction.

### Cross-context and transfer modeling

Generalization across biological conditions or cell types has attracted growing interest. Domain adaptation frameworks such as scArches (Lotfollahi et al. 2020), CSLAN (Wu et al. 2025) and scVI (Lopez et al. 2018) learn shared latent spaces to transfer representations between datasets. Cross-species or cross-tissue prediction has been explored using contrastive or attention-based models (Cao et al. 2022; Chen et al. 2022). However, most prior work focuses on embedding alignment or zero-shot annotation rather than time-evolving dynamics. Our formulation instead quantifies and predicts consistent temporal behavior across cell types through the proposed Dynamic Consistency Index (DCI) and uncertainty-aware recurrent modeling.

## Dynamic Consistency Index (DCI)

Temporal modeling in heterogeneous single-cell datasets requires identifying genes whose expression changes follow reproducible trends across cell types. Some genes display monotonic activation or suppression following injury, while others fluctuate idiosyncratically or remain nearly constant.

To quantify this difference, we introduce the **Dynamic Consistency Index (DCI)**, a simple scalar measure that captures the degree to which a gene’s temporal trajectory is aligned across cell types.

### Definition

For a given gene *g*, let *µ*_*c,t*_ denote its mean expression for cell type *c* at timepoint *t*. Take our trauma dataset as an example, which contains four timepoints representing the progression of immune response after injury: *Ctrl, <*4 h, 24 h, and 72 h. We compute the logarithmic temporal change vector

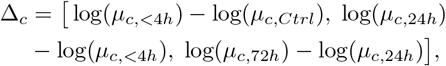

where the difference is computed using temporal differences in the logarithm of mean expression, Δ log(*µ*_*t*_) = log(*µ*_*t*+1_) − log(*µ*_*t*_), rather than raw differences. The logarithmic scale converts multiplicative fold changes in expression into additive increments, allowing cosine similarity to measure alignment of temporal *direction* rather than magnitude. It also stabilizes variance across genes that differ by orders of magnitude in expression level. Because some genes or cell types may have near-zero mean expression, we add a small constant *ε* (e.g. 10^−3^) before taking the logarithm, computing log(*µ* + *ε*) to avoid undefined values and to model a minimal background expression level. This “soft” log-transform preserves relative dynamics for highly expressed genes while preventing numerical instability for lowly expressed ones. This three-dimensional vector summarizes the directional pattern of change over the experimental timeline for each cell type. Let 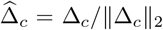 be its normalized form. The pairwise alignment between two cell types *c*_*i*_ and *c*_*j*_ is measured by cosine similarity

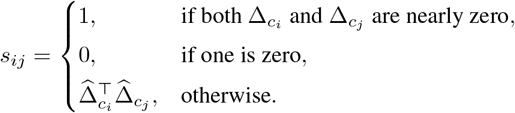

The *Dynamic Consistency Index* of gene *g* is the average of all pairwise similarities across *C* cell types:

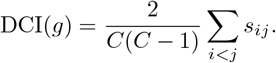

By construction DCI(*g*) ∈ [−1, 1], with higher values indicating stronger cross-cell-type coherence in temporal direction.

### Interpretation

A high DCI implies that different cell types undergo expression changes of similar sign and relative magnitude, suggesting a coordinated transcriptional program or shared regulatory control. Low or negative DCI indicates inconsistent or opposing trends, reflecting either divergent regulation or measurement noise. In practice, we treat genes with DCI ≥ 0.8 as *dynamically consistent* and model them individually over time. Genes with low DCI are excluded since their temporal profiles provide little predictive signal beyond random fluctuation.

### Properties

DCI is insensitive to global scaling of expression levels and depends only on the *direction* of change in log space, which stabilizes comparisons across genes with different expression magnitudes. The index increases monotonically with the average pairwise cosine alignment, making it interpretable as a measure of temporal coherence analogous to a correlation coefficient. Empirically, DCI correlates strongly with model predictability: genes with higher DCI yield lower mean absolute error (MAE) under all modeling frameworks. This observation supports the hypothesis that temporal consistency, rather than mere variance, determines whether a gene’s dynamics can be reliably learned.

### Practical use

Before model training, DCI is computed for every gene using the aggregated cell-type summaries. High-DCI genes define a subset with interpretable temporal structure, forming the input space for subsequent Gaussian recurrent modeling described in the next Section. This filtering step removes noise-dominated trajectories and reduces sample imbalance across cell types, improving both convergence and generalization. In addition, during the modeling phase, DCI is further incorporated into the loss function through a DCI-alignment regularization term, guiding the recurrent network to capture characteristic fold-change dynamics that remain consistent across cell types. This joint use of DCI—in both gene selection and model optimization—encourages the network to learn biologically coherent temporal patterns rather than overfitting to cell-type–specific fluctuations.

## Uncertainty-Aware Recurrent Modeling

Modeling temporal expression from aggregated scRNA-seq data requires representations that capture sequential structure while handling heterogeneity across cell types. Expression trajectories following trauma are often smooth but differ in amplitude or timing among cell types, leading to uneven noise levels and sample sizes. A deterministic regression model trained on pooled data risks overfitting dominant cell types or memorizing their mean profiles. To address this, we adopt a recurrent neural network trained under a Gaussian negative log-likelihood (NLL) objective, which learns both the expected temporal trend and its predictive uncertainty while maintaining generalization across cell types.

### Problem formulation

For a given gene *g*, let *x*_*c,t*_ ∈ ℝ_*d*_ denote its feature vector summarizing expression statistics in cell type *c* at time *t*, and *y*_*c,t*+1_ the corresponding mean expression at the next timepoint. Each training instance consists of (*x*_*c,t*− 2:*t*_, *y*_*c,t*+1_), representing three consecutive observations followed by the next-time target. The model learns a function

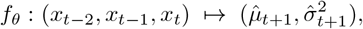

where 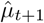 is the predicted next-time mean and 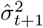 its estimated variance.

To evaluate genuine generalization, training and testing are performed on *disjoint sets of cell types*. This cross-cell-type learning setup prevents information leakage from overlapping temporal profiles and reflects the practical goal of predicting temporal behavior in unseen or sparsely sampled cell populations. It also serves as a stringent test of model inductive bias: a well-calibrated temporal model should capture consistent dynamics that transfer across biological contexts, not merely memorize cell-type-specific baselines.

### Gaussian NLL loss

For a collection of *N* training samples {(*x*_*i*_, *y*_*i*_, *s*_*i*_)}, where 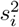 represents label-side uncertainty (empirical variance of the mean), we minimize

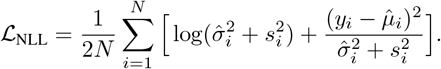

This heteroscedastic objective down-weights uncertain observations and encourages calibrated predictive variances. Unlike deterministic losses, it explicitly distinguishes epistemic error from aleatoric noise—an essential property for heterogeneous scRNA-seq data.

### DCI Alignment Loss

To incorporate cross-cell-type dynamic consistency into model training, we introduce a DCI-based regularization term that aligns each predicted temporal change with the characteristic dynamics observed in the training cell types. For each gene *g*, let 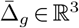 denote the mean log-difference vector across all training cell types:

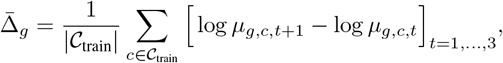

which represents the consensus fold-change trajectory of gene *g* from Ctrl → *<*4h → 24h → 72h. During training, for each predicted sequence we compute the predicted log-delta 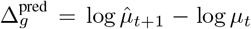 and penalize its deviation from 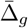 via

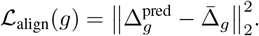

The overall training objective combines this alignment penalty with the heteroscedastic Gaussian negative log-likelihood (NLL):

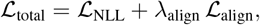

where *λ*_align_ controls the strength of the DCI prior. This encourages the model to learn temporal transitions that are not only statistically accurate but also consistent with biologically reproducible fold-change patterns observed across cell types.

### Architecture Overview

We use a gated recurrent unit (GRU) with hidden size *h* = 64 to encode temporal dependencies. Two linear heads map the hidden state to the predicted mean and log-variance, respectively, with a softplus activation ensuring non-negative variance outputs. Dropout (*p* = 0.1) is applied after each recurrent step for regularization. All models are trained using Adam (learning rate 10^−3^, weight decay 10^−4^) with early stopping based on validation performance. Identical preprocessing and hyperparameters are used across genes for fair comparison.

### GRU Network

At each step, the GRU receives a feature vector summarizing the current cell-type statistics (e.g., mean, variance, fraction of positive cells) together with DCI-derived priors, and updates its hidden state to capture dynamic patterns across timepoints. The GRU outputs both the predicted mean expression 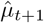 and a corresponding variance 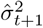, which parameterize a heteroscedastic Gaussian distribution for the next-time prediction. This architecture enables the model to learn nonlinear temporal relationships while jointly estimating predictive uncertainty, forming the backbone of our Gaussian NLL and DCI-regularized training objective.

### Interpretation

The recurrent structure enforces smooth propagation of temporal information while remaining flexible to local fluctuations. By jointly predicting mean and variance, the model captures both systematic temporal patterns and context-specific uncertainty. Cross-cell-type training emphasizes generalizable trends that persist beyond individual lineages, aligning with the biological objective of identifying reproducible temporal responses.

## Experiments and Results

### Dataset and Preprocessing

The dataset we use is from (Chen et al. 2021), which provides longitudinal single-cell RNA-seq measurements from human trauma patients (and matching control samples) collected at four timepoints: Ctrl, *<*4 h, 24 h, and 72 h. The published data include both raw single-cell profiles and aggregate per-sample gene expression summaries. In our analysis, we group cells by annotated cell types and compute per-gene summary statistics for each cell type at each time-point.

From this processed data, we construct a task of predicting next-timepoint mean expression per gene per cell type, using only the trained subset of cell types to learn dynamics and evaluating on disjoint, held-out cell types. This cross-cell-type prediction framework, grounded in real trauma scRNA-seq data, tests whether learned temporal models generalize across heterogeneous cellular contexts.

Cells are annotated into multiple immune cell types, and expression is aggregated at the cell-type level to form a compact temporal representation. For each gene and cell type, we compute ten summary statistics—mean (*µ*), standard deviation, fraction of positive cells, mean of positives, median, interquartile range, 10th and 90th quantiles, sample count, and standard error. These statistics constitute the feature vector at each timepoint.

All expression values are log-transformed to stabilize variance. We compute the Dynamic Consistency Index (DCI) for every gene as described previously, and select those with DCI ≥ 0.8 for main experiments. Low-DCI genes are excluded because their irregular dynamics provide no stable temporal signal. For each experimental run, we randomly sample 10 high-DCI genes and train the model on this subset; we repeat this procedure for 10 independent trials and report the average performance across runs. No normalization across cell types is applied beyond the use of standardized features, since relative differences are biologically meaningful.

### Feature Construction and Train/Test Split

Each training sample corresponds to a single gene in one cell type, represented by its temporal summary statistics over three consecutive timepoints (*t* − 2, *t* − 1, *t*) and the target mean expression at *t*+1. This produces short time sequences of fixed length suitable for recurrent or feedforward models.

To evaluate generalization across biological contexts, we adopt a cross-cell-type split: cell types are partitioned into disjoint train, validation, and test sets. All genes are trained on the same cell-type partition to avoid leakage. This setup ensures that no cell-type-specific expression profile seen during training appears in testing. It mimics the practical scenario where models are trained on well-sampled cell populations but applied to unseen or sparsely measured ones.

### Baselines

We compare the proposed recurrent Gaussian model with several deterministic and probabilistic baselines:

- **Naїve Carry-Forward:** predicts 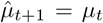, serving as a lower bound.
- **Linear predictor:** fits 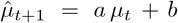 using least-squares on training cell types.
- **MLP:** a two-layer feedforward network trained with L1 loss.
- **MLP + Gaussian NLL:** identical architecture with heteroscedastic output trained under Gaussian NLL.
- **Transformer:** a small encoder with self-attention and Gaussian NLL loss.
- **RNN:** a gated recurrent unit trained with L1 loss.
- **RNN + Gaussian NLL:** the proposed uncertainty-aware recurrent model.

All models share identical preprocessing and feature dimensions, differing only in temporal parameterization and loss function.

### Training Details

Each model is trained independently for every gene. Features are standardized within the training set, and identical hyperparameters are used across baselines for fairness. The recurrent models use a GRU with hidden size 64, dropout rate 0.1, and two linear output heads for mean and log-variance. The MLP consists of two fully connected layers (128 and 64 units) with ReLU activation and dropout. Optimization uses Adam with learning rate 10^−3^ and weight decay 10^−4^. Early stopping is based on validation MAE with patience of 20 epochs. All experiments are repeated with three random cell-type splits, and results are averaged.

### Evaluation Metrics

We report the following metrics for each gene and model:

- **MAE:** mean absolute error between predicted and observed mean expression.
- **95% Coverage:** For models with Gaussian NLL, empirical proportion of test samples whose true value lies within the predicted 95% confidence interval.

For clarity, we report per-gene MAE values and aggregate means across all test genes. Lower MAE and NLL, and coverage close to 95%, indicate better accuracy and calibration.

### Key Observations

As shown in table 1, high-DCI genes exhibit predictable temporal trends, while low-DCI genes show near-random behavior. The RNN with Gaussian NLL consistently achieves the lowest MAE and best uncertainty calibration across test cell types. Gaussian NLL reduces overconfidence and stabilizes training compared to L1 loss, especially for noisy cell types. The linear drift baseline performs competitively on monotonic genes but fails to capture nonlinear recovery patterns. These results confirm that combining DCI-based gene selection with uncertainty-aware recurrent modeling yields robust and interpretable temporal predictions in heterogeneous single-cell systems.

**Table 1:** MAE scores on test cell types.

|  | Example Genes |  |  |  |  | Mean | 95%coverage |
| --- | --- | --- | --- | --- | --- | --- | --- |
|  | GCA | TRAV22 | HBA1 | ATM | AREG |  |  |
| Naïve | **0.0725** | 0.0005 | 0.98 | 0.4352 | 0.4015 | 0.2765 | / |
| Linear | 0.4755 | **0.0004** | 2.352 | 0.3789 | 0.5202 | 0.485 | / |
| MLP | 0.1103 | 0.0351 | 0.65 | 0.3346 | 0.2785 | 0.2752 | / |
| MLP+Gaussian | 0.1956 | 0.0115 | 0.6339 | 0.343 | 0.3138 | 0.2582 | 0.889 |
| Transformer | 0.119 | 0.0064 | 0.6495 | 0.3901 | 0.2778 | 0.2457 | / |
| GRU | 0.0973 | 0.0057 | 0.655 | 0.4022 | **0.2559** | 0.2369 | / |
| GRU+Gaussian | 0.128 | 0.0029 | **0.5911** | **0.306** | 0.2613 | **0.2229** | 0.911 |

### Role of the Dynamic Consistency Index (DCI)

#### Visual intuition on TRAIN

We first illustrate how DCI captures cross–cell-type temporal regularity. For a small panel of genes, we overlay cell-type mean trajectories (*µ* over *Ctrl* → *<4h* → *24h* → *72h*) using only the *training* cell types. High-DCI genes exhibit aligned directional changes across cell types, whereas low-DCI genes show heterogeneous or opposing trends. These plots provide an immediate, leakage-free view that DCI reflects coherent temporal structure rather than scale alone. (See Fig. 2.)

**Figure 1:**
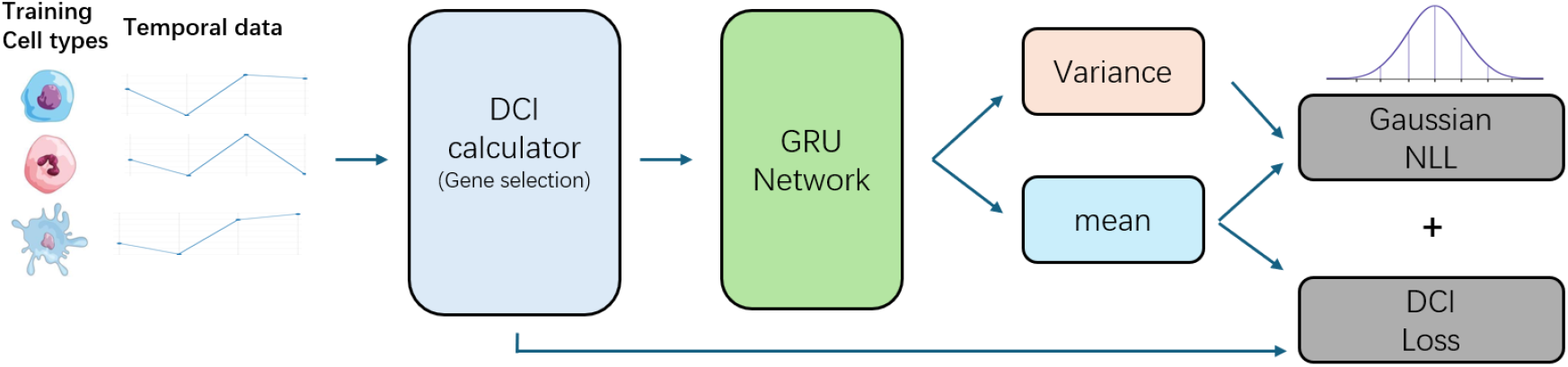
Overview of the workflow. Temporal scRNA-seq data from training cell types are used to compute the Dynamic Consistency Index (DCI), which quantifies reproducible fold-change patterns across time and guides gene selection. The resulting DCI priors are fed into the GRU network along with cell-type summary statistics. The network outputs both the predicted mean and variance of gene expression at the next timepoint. Training minimizes a heteroscedastic Gaussian negative log-likelihood (NLL) combined with a DCI alignment loss, encouraging predictions that remain consistent with characteristic fold-change dynamics observed across cell types.

**Figure 2:**
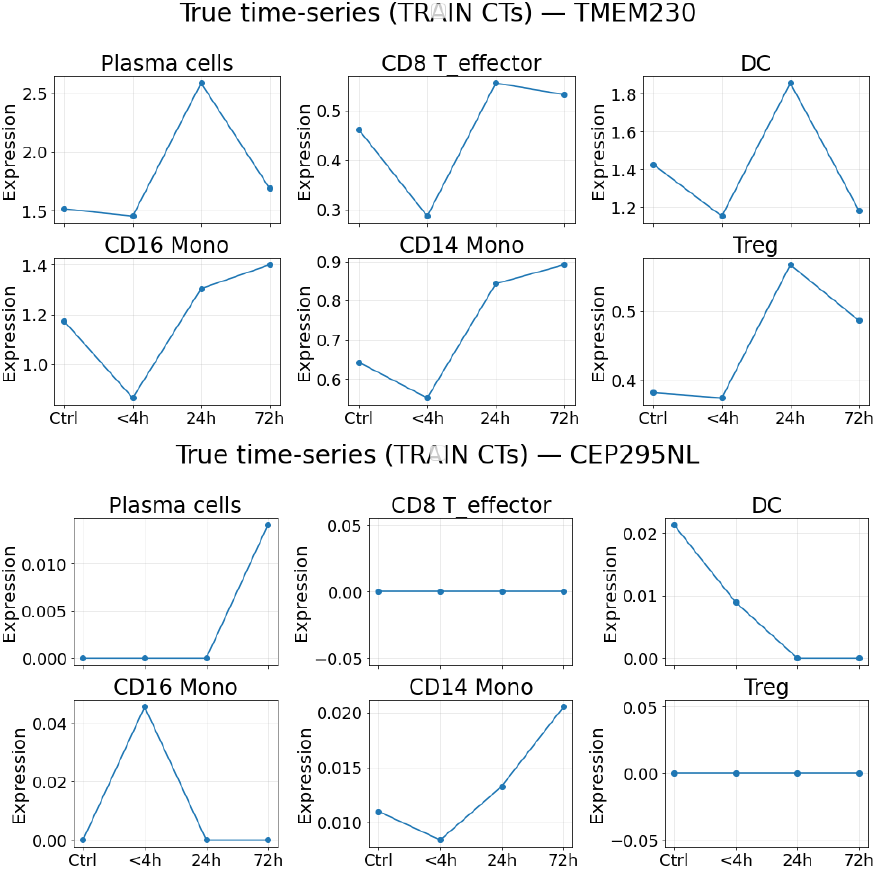
Visual intuition for the Dynamic Consistency Index (DCI). Each panel shows temporal expression trajectories (*µ*) across sampled training cell types for a single gene, from *Ctrl* to *72h*. (**Top 6 panels**) High-DCI gene (DCI=0.84): cell-type trajectories exhibit coherent, aligned directional changes, reflecting a shared temporal trend. (**Bottom 6 panels**) Low-DCI gene (DCI=-0.02): trajectories vary in direction and amplitude, indicating inconsistent or noise-dominated behavior. These examples demonstrate that DCI effectively distinguishes genes with reproducible temporal dynamics from those with irregular patterns.

### Generalization of DCI across cell types

To examine whether the temporal regularity captured by DCI is stable beyond the training cell types, we compute DCI_train_ using only the training cell types and DCI_all_ using both training and test cell types. This analysis evaluates whether temporal directionality identified in well-sampled populations persists when new, unseen cell types are added to the cohort.

Figure 3 shows the scatter of DCI_train_ versus DCI_all_ for all genes. The two measures exhibit a strong and nearly linear relationship (Spearman *ρ* = 0.923, Pearson *r* = 0.933, *p <* 10^−15^). An ordinary least-squares fit yields a slope of 0.81 and an intercept of 0.02, indicating that the overall DCI values remain high when additional cell types are included, with only a modest downward adjustment due to increased biological variability. Most points cluster tightly along the identity line, showing that genes with consistent temporal dynamics in training retain similar coherence when unseen cell types are introduced.

**Figure 3:**
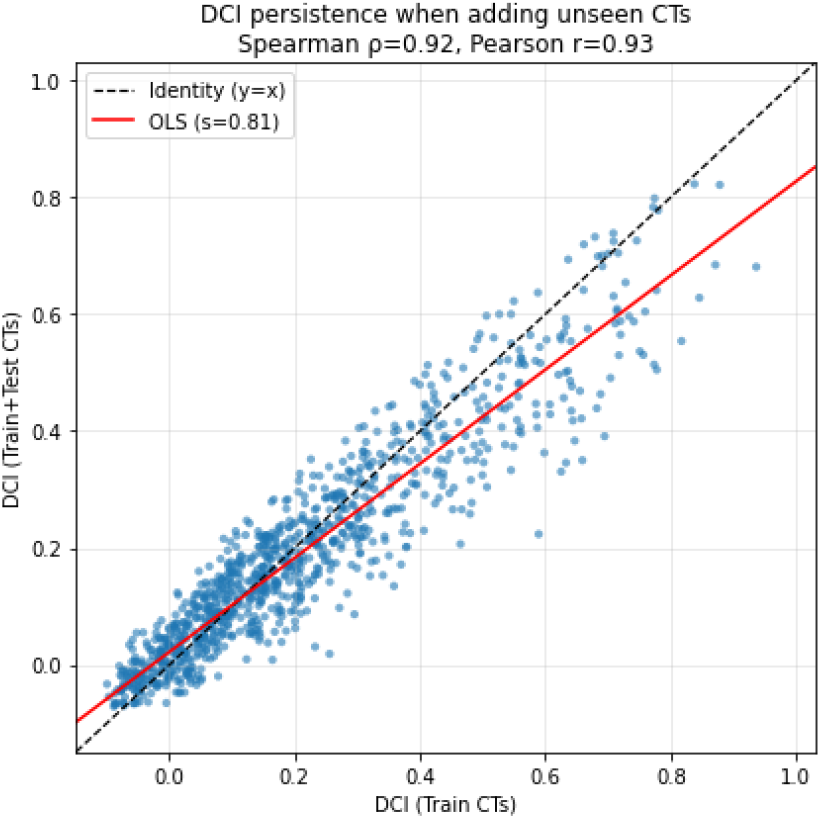
Generalization of DCI across cell types. Scatter plot of DCI_train_ versus DCI_all_ showing strong agreement (Spearman *ρ* = 0.923, Pearson *r* = 0.933). The dashed black line indicates the identity (*y* = *x*), and the red line denotes the OLS fit (slope 0.81, intercept 0.02). Each point represents a single gene.

These results demonstrate that DCI reflects an intrinsic property of a gene’s temporal behavior rather than a partition-specific artifact. High-DCI genes remain dynamically consistent across broader cell-type contexts, confirming that the DCI metric generalizes robustly and can be used as a reliable indicator of predictability and biological stability.

### Predictability as a Function of DCI

To quantify how dynamic consistency influences cross-cell-type generalization, we stratify genes by their training Dynamic Consistency Index (DCI) into five bins: [0, 0.2), [0.2, 0.4), [0.4, 0.6), [0.6, 0.8), and [0.8, 1.0]. Within each bin, we evaluate all models on held-out cell types and report mean test performance.

#### Relative error metric

While MAE gives an absolute error measure, it does not directly capture how much better a model is compared to a trivial baseline. We therefore adopt the *mean absolute scaled error* (MASE), which was introduced by Hyndman & Koehler (2006) as a scale-independent benchmarked error metric in forecasting literature (Hyndman and Koehler 2006). MASE is defined for each gene *g* as

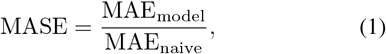

where MAE_naive_ is the test error of a carry-forward predictor 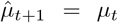. MASE values below 1 indicate that the model improves upon the naїve baseline, while values above 1 suggest that temporal dynamics are too inconsistent for the model to outperform a simple persistence assumption.

#### Empirical trends

Figure 4 plots MASE across DCI bins for all models. We observe a clear monotonic relationship: as DCI increases, MASE consistently decreases, indicating that genes with more consistent temporal patterns are easier to predict. In the lowest-DCI bin ([0, 0.2)), all models exhibit MASE *>* 1, implying that even complex architectures cannot improve upon the naїve baseline when gene dynamics are erratic. In contrast, for highly consistent genes (DCI *>* 0.8), our heteroscedastic recurrent model (RNN+Gaussian NLL) achieves the lowest MASE (0.78), corresponding to a 22% reduction in MAE relative to the naїve baseline. Transformer and MLP baselines show similar downward trends but saturate around MASE≈ 0.85.

**Figure 4:**
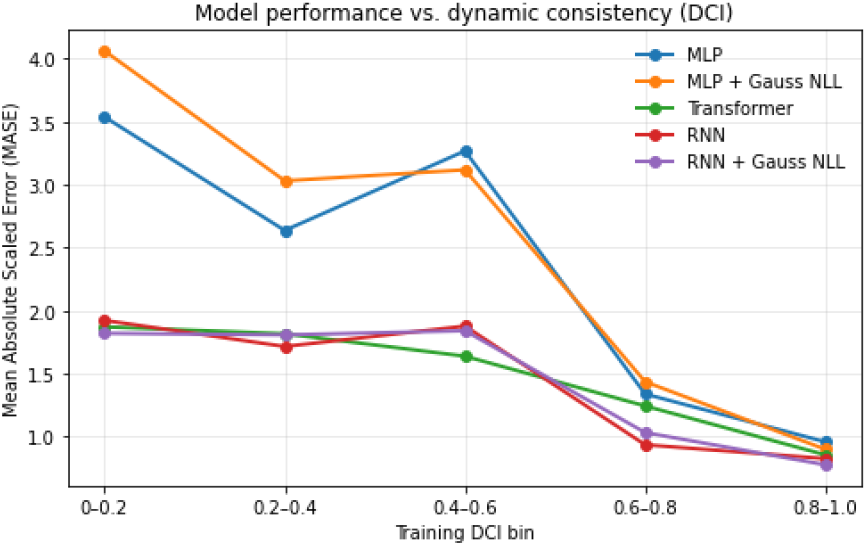
Predictability as a function of dynamic consistency. Mean Absolute Scaled Error (MASE) across DCI bins [0, 0.2), [0.2, 0.4), [0.4, 0.6), [0.6, 0.8), and [0.8, 1.0]. Lower MASE values indicate improved performance relative to a naїve baseline. As DCI increases, MASE consistently decreases across all models, demonstrating that genes with more consistent temporal dynamics are easier to predict. The RNN+Gaussian NLL model achieves the lowest MASE at high DCI, confirming its superior cross-cell-type generalization.

#### Interpretation

These results confirm that the Dynamic Consistency Index captures a meaningful notion of predictability: higher DCI implies more reproducible temporal evolution across cell types and, correspondingly, lower normalized prediction error. MASE provides a complementary scale-free view of generalization—showing not just that RNN+Gaussian NLL yields the lowest absolute MAE, but that it also provides the largest relative gain over a naїve carry-forward predictor. Together, DCI and MASE jointly quantify the alignment between biological regularity and model generalization capacity.

#### Leakage control and robustness

All DCI values used for selection or binning are computed on training cell types only; test cell types are held out throughout.

Train/validation/test partitions, preprocessing, and early-stopping criteria are identical across models. We repeat the DCI–predictability analysis across several random partitions and report aggregated statistics to ensure that conclusions are not split-specific.

#### Takeaway

These studies establish DCI as (i) a visual and quantitative marker of coherent temporal dynamics across cell types, (ii) a property that generalizes to unseen cell types, and (iii) a practical predictor of downstream modeling difficulty, with the uncertainty-aware recurrent model performing best in the high-DCI regime.

## Conclusion

In this work, we study temporal modeling of gene expression across cell types in human trauma scRNA-seq data. We introduce the Dynamic Consistency Index (DCI) to quantify reproducible temporal trends and demonstrate its strong correlation with cross-cell-type predictability. By combining DCI-based gene selection with a recurrent model trained under a Gaussian negative log-likelihood objective, we obtain improved accuracy and calibrated uncertainty compared to deterministic baselines.

Our analysis shows that high-DCI genes display coherent temporal behavior that generalizes well to unseen cell types, while low-DCI genes remain inherently unpredictable. These findings suggest that temporal consistency, rather than variance magnitude, is the key determinant of model learnability in single-cell dynamics.

Beyond trauma, the proposed DCI framework can generalize to other longitudinal single-cell or perturbation datasets, such as infection, drug response, or aging studies. Future work could extend the current formulation by integrating neighborhood information among genes or pathways, enabling graph-based priors that capture co-regulatory structure. Another promising direction is coupling DCI with latent dynamic models or diffusion frameworks to predict full-trajectory evolution rather than next-time averages. These extensions would broaden the applicability of DCI-guided temporal modeling and deepen its connection with systems-level biological interpretation.

We expect the proposed DCI framework and uncertainty-aware modeling strategy to serve as a foundation for studying temporal regularities in other longitudinal or cross-condition single-cell datasets.

## Data Availability

The datasets and code used in this study are available upon reasonable request. A public release is planned and will be shared via an online repository upon publication.

## Acknowledgments

This section will be included in the final version.

